# Estimating mutual information under measurement error

**DOI:** 10.1101/852384

**Authors:** Cong Ma, Carl Kingsford

## Abstract

Mutual information is widely used to characterize dependence between biological signals, such as co-expression between genes or co-evolution between amino acids. However, measurement error of the biological signals is rarely considered in estimating mutual information. Measurement error is widespread and non-negligible in some cases. As a result, the distribution of the signals is blurred, and the mutual information may be biased when estimated using the blurred measurements. We derive a corrected estimator for mutual information that accounts for the distribution of measurement error. Our corrected estimator is based on the correction of the probability mass function (PMF) or probability density function (PDF, based on kernel density estimation). We prove that the corrected estimator is asymptotically unbiased in the (semi-) discrete case when the distribution of measurement error is known. We show that it reduces the estimation bias in the continuous case under certain assumptions. On simulated data, our corrected estimator leads to a more accurate estimation for mutual information when the sample size is not the limiting factor for estimating PMF or PDF accurately. We compare the uncorrected and corrected estimator on the gene expression data of TCGA breast cancer samples and show a difference in both the value and the ranking of estimated mutual information between the two estimators.

## 1 Introduction

Mutual information is an important measure to evaluate the degree of dependence between biological signals. It compares the joint probability of two signals with their marginal probability and is able to capture both linear and non-linear dependencies. However, when the biological signals are measured with error, the error blurs the probability distribution of the signals and leads to inaccurate mutual information estimates. Current studies on mutual information, both from a theoretical perspective and from an application perspective, mostly assume the observed signals are accurate. Ignoring the potential measurement error leads to biases in the estimated mutual information between the signals.

Applications of mutual information in computational biology include analyzing the co-evolution relationship between amino acids or nucleotides [1], inferring co-occurrence patterns of protein domains [2], constructing gene regulatory networks [3], studying neural connectivity circuits [4], and so on. An accurate estimation of mutual information is critical to these studies.

Measurement error for some biological signals is non-negligible, especially when high resolution of the measurement is needed. For example, inferring transcript-level abundances tends to be more error-prone than inferring gene-level abundances [5]. However, transcript-level abundances are able to reveal more detailed changes between samples such as differential isoform usage [6]. Recent single-cell measurements [7–12] suffer from more measurement error, of which the causes include doublets [13] and dropouts [14]. The increasing degree of measurement error poses a challenge of accurately estimating the mutual information of the true biological signals. In general, the high noise-to-signal ratio is challenging for revealing patterns in many fields of analyses.

Measurement error has been modeled and used in analyses in some areas, but it is not known how to incorporate these errors into the mutual information estimation. For example, a correction term for measurement error has been developed for Pearson correlation by Spearman [15]. In expression quantification, the measurement error has been modeled and integrated into the analysis of detecting differentially expressed genes/transcripts. Some quantification methods [16, 17] use bootstrapping or Gibbs sampling strategies to estimate a series of possible abundances to represent the scale of the measurement error. As shown by Pimentel et al. [18] and Zhu et al. [19], incorporating a model of measurement error into the differential expression detection methods improves the accuracy. This inspires the incorporation of the measurement error in other analyses.

Many theoretical studies about mutual information focus on correcting biases due to small sample sizes or deriving better estimation for probability density function (PDF). However, error-free measurements are usually implicitly assumed in these studies. Basharin [20] focused on discrete distributions and derived a correction term for the estimation bias due to small sample sizes in estimating probability mass function (PMF), which is further used in mutual information calculation. Moon et al. [21] proposed a mutual information estimator when the random variables follow continuous distributions. Their estimator uses kernel density estimation (KDE) with the bandwidth suggested in Silverman [22] to estimate PDF. Kraskov et al. [23] later proposed a *k*-nearest neighbor (KNN) approach for estimating PDF in mutual information calculation. Khan et al. [24] compared KNN-based mutual information estimator and the KDE-based one and characterized the cases where one is better than the other. Their comparison includes the case where signals are measured with error, but adaptation or correction for the error is not proposed. Holmes and Nemenman [25] based their work on the KNN mutual information estimator and presented a strategy to use bootstrapping of samples to estimate the error bars of estimated mutual information. Zeng et al. [26] developed a mutual information estimator based on copula density estimation [27] with the Jackknife approach.

In this work, we derive a corrected mutual information estimator that reduces the inaccuracy caused by measurement error. Our corrected estimator is based on a correction for the estimated PMF in the (semi-) discrete case or for the KDE in the continuous case using the distribution of measurement error. We prove that in the (semi-) discrete case our corrected estimator is asymptotically unbiased when the measurement error distribution is known. We discuss the assumptions under which the corrected estimator in the continuous case leads to a reduced bias in mutual information estimation. Using simulated data, we show that our corrected estimator is more accurate than the baseline estimator that plugs in the average bootstraps of observations in the (semi-) discrete or KDE-based mutual information estimators. The result shows that the derivation is correct and our corrected estimator effectively reduces the bias of mutual information estimation. Using the corrected and uncorrected estimator to calculate the pairwise mutual information between gene expression estimates on 1168 breast cancer samples in The Cancer Genome Atlas (TCGA, https://cancergenome.nih.gov), we observe a high Spearman correlation between the results of both estimators in general. However, the specific values of estimated mutual information vary as much as 40%, and the sets of genes with high estimated mutual information differ. Therefore, when the values of mutual information are of interest or the ranking of the subset of genes with the highest mutual information is needed, the effect of measurement error cannot be ignored.

## 2 Methods

### 2.1 Problem setup and baseline solution

Mutual information *I*(*X, Y*) between random variables *X* and *Y* is defined by the Kullback–Leibler divergence between the joint distribution and the multiplication of its two marginal distributions:

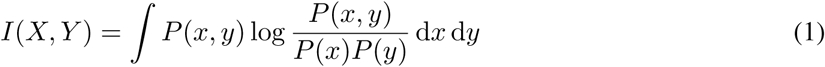

where *P* (*x, y*) is the joint probability density or probability mass function, *P* (*x*) and *P* (*y*) are the marginal probability density or mass functions of random variables *X* and *Y*. *P* (*x, y*) is usually not known and is estimated from samples. Mutual information captures the probability dependence between two random variables, specifically, whether the value of one random variable changes the belief of what value the other random variable takes.

In the presence of measurement error, we assume the true values and observed values of two biological signals (*ξ*_*x*_ and *ξ*_*y*_) are generated as follows. The joint distribution 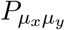 describes the likelihood of true values of signals *ξ*_*x*_ and *ξ*_*y*_ across all samples. Let (*µ*_*xs*_, *µ*_*ys*_) be the true intensities of the two signals in sample *s*. (*µ*_*xs*_, *µ*_*ys*_) is drawn from the distribution 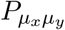. The measurement error follows the distribution 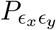 and assumed to affect the true signals through addition. Measurement errors of signal *ξ*_*x*_ and of *ξ*_*y*_ may affect each other, and thus we do not assume the independence between *ϵ*_*x*_ and *ϵ*_*y*_. Suppose the true signal (*µ*_*xs*_, *µ*_*ys*_) of sample *s* is measured multiple times and let (*ϵ*_*xsj*_, *ϵ*_*ysj*_) be the error of the *j*^*th*^ measurement, the *j*^*th*^ observation is

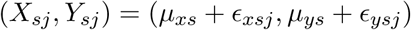

Given the observation (*X*_*sj*_, *Y*_*sj*_) of *S* samples each measured *B* times, we would like to estimate the dependence between the true signals *I*(*µ*_*x*_, *µ1*_*y*_).

Treating the observed signal values (*X*_*sj*_, *Y*_*sj*_) as the true values (*µ*_*xs*_, *µ*_*ys*_) leads to an inaccurate estimation of mutual information *I*(*µ*_*x*_, *µ*_*y*_). Because the probability density or mass function of the observation is different from that of the true signal values. Let *P*_*xy*_ be the probability density or mass function of the observation. It has the following relationship with the distribution of the true signal values:

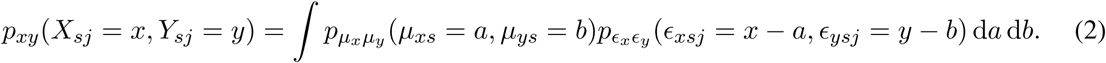

Similarly, the marginal distributions of *X*_*sj*_ and *Y*_*sj*_ are different from that of *µ*_*xs*_ and *µ*_*ys*_. Since *P*_*xy*_, *P*_*x*_, and *P*_*y*_ differ from *P*_*µxµy*_, 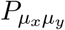, and *P*_*µy*_, the mutual information calculated using the distribution of the observation also differs from *I*(*µ*_*x*_, *µ*_*y*_).

Our goal is to derive an accurate estimation of the mutual information between *µ*_*x*_ and *µ*_*y*_ using the observations *X*_*sj*_ and *Y*_*sj*_. We measure the accuracy by whether the bias of estimation is close to zero, where the bias is the difference between the expectation of the estimated and the true mutual information.

We further assume that the measurement error has zero expectation and that follows the same distribution across all samples. Based on these assumptions, we derive a corrected mutual information estimator for both the semi-discrete case and the continuous case separately. Finally, we relax the assumption that the error distribution is the same and extend our corrected estimators.

### 2.2 Semi-discrete case: correcting the probability mass function using a transition matrix

In the semi-discrete case, we assume that both the true signals (*µ*_*x*_, *µ*_*y*_) and the observed signals (*X, Y*) are real-valued. Observed signals (*X, Y*) are a perturbation of (*µ*_*x*_, *µ*_*y*_) by adding measurement errors. The real-valued space is partitioned into several categories. Let 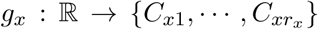 be the category mapping function that maps the real-valued signal of *ξ*_*x*_ to its corresponding categories, and 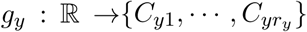 be the category mapping function for signal *ξ*_*y*_. The two category mapping functions can be combined by *g*: ℝ ℝ *C*_1_, …, *C*_*n*_, which is defined as *g*(*a, b*) = (*g*_*x*_(*a*), *g*_*y*_(*b*)) for any signal intensity *a* of *ξ*_*x*_ and intensity *b* of *ξ*_*y*_, and the pairs of categories of (*C*_*xj*_, *C*_*yk*_) are reparameterized using *C*_*i*_. *g* informs the categories of both *ξ*_*x*_ and *ξ*_*y*_ simultaneously, and the probability mass function (PMF) of *g*(*X, Y*) (or *g*(*µ*_*x*_, *µ*_*y*_)) informs both the PMF of *g*_*x*_(*X*) (or *g*_*x*_(*µ*_*x*_)) and the PMF of *g*_*y*_(*Y*) (or *g*_*y*_(*µ*_*y*_)). The mutual information between the categories, *g*_*x*_(*µ*_*x*_) and *g*_*y*_(*µ*_*y*_), is of interest. For example, *µ*_*x*_ and *µ*_*y*_ are the probability of a logistic regression that has two categories, high and low (Figure 1). *X* and *Y* are estimated probabilities, which may fall into a different category from the true signals. We assume that *g* is known, which is a reasonable assumption when the partition of the real-valued space is biologically meaningful. Our goal is to derive an estimator for the mutual information between *g*_*x*_(*µ*_*x*_) and *g*_*y*_(*µ*_*y*_) using the observation *g*_*x*_(*X*) and *g*_*y*_(*Y*).

**Figure 1:**
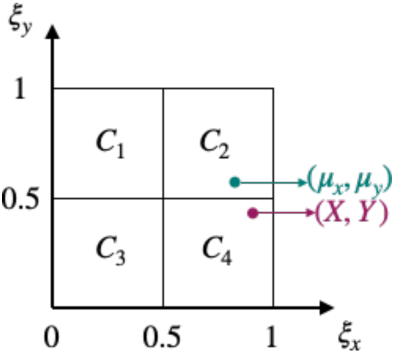
Example of real-valued (*µ*_*x*_, *µ*_*y*_) and the partition of the space into four categories. Observation (*X, Y*) is different from (*µ*_*x*_, *µ*_*y*_) due to measurement error. Here *g*(*µ*_*x*_, *µ*_*y*_) = *C*_2_ and *g*(*X, Y*) = *C*_4_.

Given sample *s* and bootstrap measure *b*, the probability of the observation *g*(*X*_*sb*_, *Y*_*sb*_) has the following relationship with *g*(*µ*_*xs*_, *µ*_*ys*_):

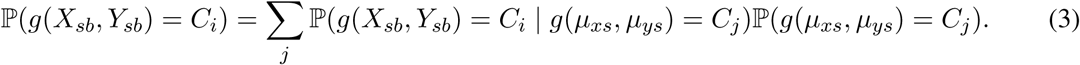

Define the following matrix *F* ∈ ℝ^*n*×*n*^, and PMF *P* ∈ ℝ^*n*^ and *Q* ∈ ℝ^*n*^:

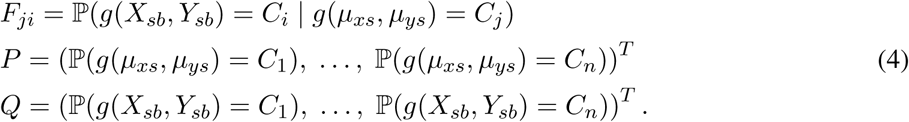

Using the above matrix and vector notations, equation (3) is equivalent to *Q*^*T*^ = *P*^*T*^*F*, where matrix *F* can be viewed as a transition matrix. We use this equality to derive the following corrected estimator.

#### Theorem 1.

*Given transition matrix F as defined in* equation (4), *measurements of signals* {*X*_*sb*_} *and* {*Y*_*sb*_} *for* 1 ≤ *s* ≤ *S and* 1 ≤ *b* ≤ *B, let* 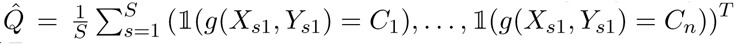, *where 𝟙 is the indicator function*. *If* 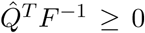 *element-wise, and the mutual information is estimated using joint PMF of* 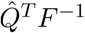 *and the corresponding marginal PMFs, then the estimated mutual information is an unbiased estimator for the target mutual information of P*^*T*^ *asymptotically as S* → ∞.

*Sketch of proof*. We use the property that mutual information is a linear combination of three entropy terms

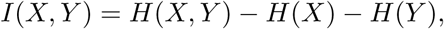

where the entropy of a (joint) discrete random variable with PMF *Q* = (*q*_1_, …, *q*_*n*_)^*T*^ is defined as

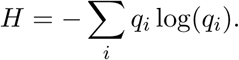

If the entropy estimator 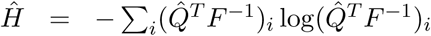 is an unbiased estimator for *H* = − Σ_*i*_ *p*_*i*_ log(*p*_*i*_), then the mutual information calculated using 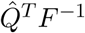 is an unbiased estimator for the mutual information calculated using *P*, where 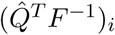 is the *i*^*th*^ element of vector 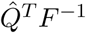.

Basharin [20] proved that entropy calculated using 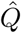 is an unbiased estimator for the entropy calculated using *Q* as the number of samples goes to infinity. We use their proof as a template and derive that the entropy of 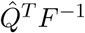 is an unbiased estimator of *Q*^*T*^ *F*^−1^ = *P* as the number of samples goes to infinity. See Lemma S2 in Appendix Section S1.1 for the details of proof that entropy of 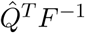 is an unbiased estimator of the entropy of *Q*^*T*^ *F*^−1^.

With a little abuse of notation, the unbiasedness also holds for the following estimator.

#### Corollary 1.1.

*Given measurements of signals* {*X*_*sb*_} *and* {*Y*_*sb*_} *for* 1 ≤ *s* ≤ *S and* 1 ≤ *b* ≤ *B*, let 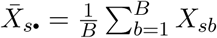 *and* 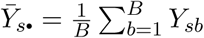. *Let transition matrix F* ∈ ℝ^*n*×*n*^ *have the following entries*

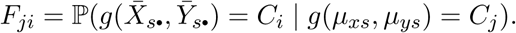

*Let the estimated PMF be* 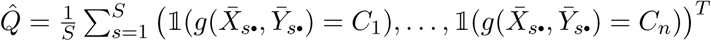, *where 𝟙 is the indicator function*. *If* 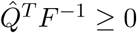 *element-wise, and the mutual information is estimated using joint PMF of* 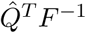 *and the corresponding marginal PMFs, then estimated mutual information is an unbiased estimator for the target mutual information of P*^*T*^ *asymptotically as S* → ∞.

The above theorem and corollary hold for both the semi-discrete case and the discrete case. However, matrix *F* is generally not known beforehand, and an estimation of *F* is required. We provide an estimation for *F* based on the real-valued assumption in the semi-discrete case.

We assume the distribution of (*µ*_*x*_, *µ*_*y*_) within each category is a uniform distribution and approximate the measurement error by a Gaussian distribution. A Gaussian distribution is appropriate to approximate the measurement error because the Central Limit Theorem states that 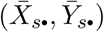 converges in distribution to a Gaussian distribution as *B* → ∞. Let 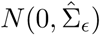 be the limiting distribution for 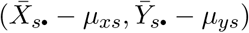. Let *U*_*i*_ = [*l*_*xi*_, *u*_*xi*_] × [*l*_*yi*_, *u*_*yi*_] be the real-valued range corresponding to *g*(*µ*_*x*_, *µ*_*y*_) = *C*_*i*_ and *U*_*j*_ = [*l*_*xj*_, *u*_*xj*_] × [*l*_*yj*_, *u*_*yj*_] be the ranges corresponding to *g*(*µ*_*x*_, *µ*_*y*_) = *C*_*j*_. The approximated transition probability is:

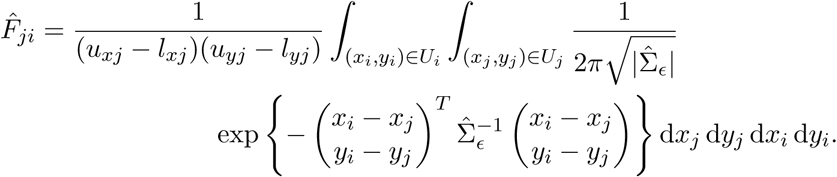

In the discrete case, correcting the distribution of (*µ*_*x*_, *µ*_*y*_) using transition matrix *F* has the same form as image denoising or deblurring [28]. Nevertheless, the goal of image deblurring is to correct the individual observation, but our estimator is based on correcting the distribution. The estimated PMF is proportional to the counts in the discrete case. The counts can be viewed as an observation from an multinomial distribution. From this perspective, correcting the counts is equivalent to correcting an multinomial observation, which explains the similarity between image deblurring and PMF correction.

### 2.3 Continuous case: correcting estimated PDF using kernel density estimation (KDE)

In this section, we aim to reduce the bias in the KDE-based mutual information estimation in the continuous case. We first introduce the formula of the KDE-based mutual information estimator and derive its estimation bias. We then derive a correction for the estimated density and prove the unbiasedness of the corrected density estimation. We finally discuss the scenarios when the corrected density reduces the error in mutual information estimation.

Moon et al. [21] develops a mutual information estimator that uses the kernel density estimation to estimate the PDF. Given true signal values (*µ*_*xs*_, *µ*_*ys*_) and using a diagonal bandwidth matrix, the kernel density estimation given by Silverman [22] is

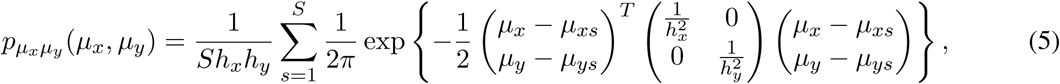

where *h*_*x*_ and *h*_*y*_ are the bandwidth for the two axes separately. The estimation of mutual information is the sample average of the differences among the log of probability densities:

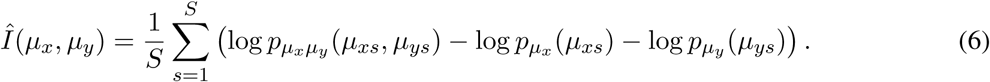

With given true values (*µ*_*xs*_, *µ*_*ys*_), these estimators are unbiased as *S* → ∞ except for the boundaries or tails of the distribution [29].

In presence of measurement error, (*µ*_*xs*_, *µ*_*ys*_) is not observed. The observation (*X*_*sj*_, *Y*_*sj*_) is a summation of the true signal and error, where the error distribution is estimable from the bootstraps. We assume the distribution of measurement error is the same for all samples. The density of the observation *P*_*xy*_ is different from the density of the true signals 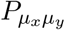 as shown by equation (2). Our goal is to derive a corrected estimator for mutual information of *I*(*µ*_*x*_, *µ*_*y*_) with a reduced bias.

The bias of each log term in mutual information estimator in equation (6) has the following upper bound.

#### Lemma 1.

*Given the true probability density p and fixed point* (*x, y*)*, for any estimator of the density at the point* 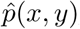, *if the true and estimated density are lower bounded by δ* > 0, *that is, p*(*x, y*) ≥ *δ and* 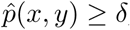, *then the bias of* 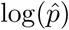 *at the point is upper bounded by*:

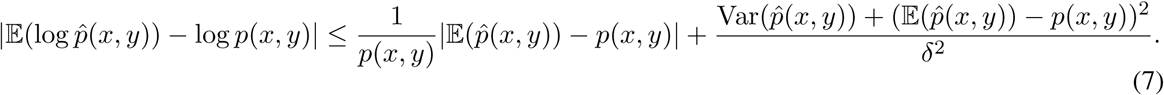

*Sketch of proof*. The three main steps of deriving the inequality are: applying Taylor expansion on the log function with mean value form of the remainder, then taking the expectation on both sides, and finally replacing the quadratic term with bias-variance decomposition. See Appendix Section S1.2 for the details.

According to this lemma, if the density estimator 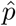 has both small bias and small variance at a nonzero density position (*x, y*), the log terms in equation (6) will have small bias. The points evaluated in equation (6) are true signal values (*µ*_*xs*_, *µ*_*ys*_), and thus the true densities and the estimated densities by KDE using the observed signals (*X*_*sj*_, *Y*_*sj*_) at these points are non-zero. We focus on reducing the bias of density estimation, 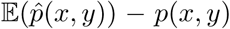, and derive the following (asymptotically) unbiased estimators for each summation term in the KDE density formula (5).

#### Theorem 2.

*Let* 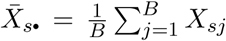 *and* 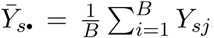. *Let W be an diagonal bandwidth matrix used in KDE*, 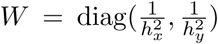. *Assuming the measurement error of* 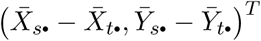 *follows a Gaussian distribution* 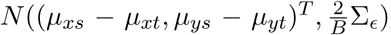, *let P and* {*ζ*_1_, *ζ*_2_} *be the eigenvectors and eigenvalues of* 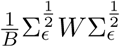. *Let* 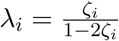, 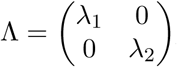, and 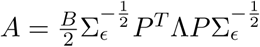. *When ζ*_*i*_ *satisfies* 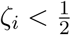 *for all i, the following estimator*,

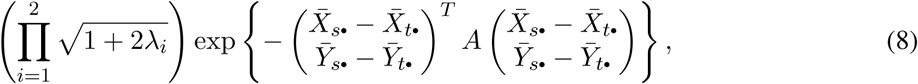

*is an unbiased estimator for the KDE term in* equation (5):

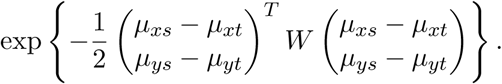

The proof of the theorem is in Appendix Section S1.3.

Given sample *s*, the covariance of error Σ_*ϵ*_ is usually not known or the error may not be a Gaussian distribution. Nevertheless, a Gaussian limiting distribution is a good approximate to model the error as shown in Central Limit Theorem:

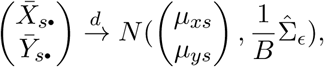

where *d* means convergence in distribution and the estimated 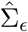 is calculated by

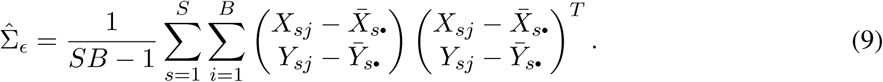

We prove that using the estimated error distribution in equation (9) leads to an asymptotically unbiased estimator for the corresponding KDE term.

#### Theorem 3.

*Let P and* {*ζ*_1_, *ζ*_2_} *be the eigenvectors and eigenvalues of* 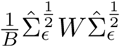. *Let*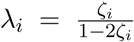, 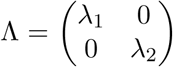, *and* 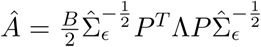. *When ζ*_*i*_ *satisfies* 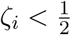 *for all i, the following estimator is an asymptotically unbiased estimator for the KDE term in* *equation* (5):

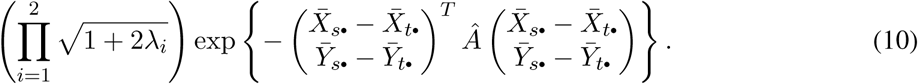

See Appendix Section S1.4 for the proof.

#### Corollary 3.1.

*Let* 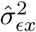 *be the variance of measurement error on signal ξ*_*x*_, *which is the top left element in covariance matrix* 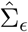. *When* 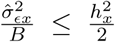, *the following estimator is an unbiased estimator for* 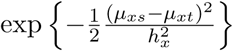:

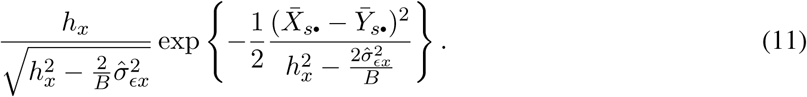

A corrected mutual information estimator can be derived by plugging in the corrected density estimator (10) and (11) into equation (6). We claim that when *λ*_*i*_ is small, the expectation of 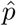 dominates the second order moment around 0, which further dominates 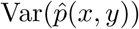.

**Claim 1.** *There exists a small positive number δ such that when* 0 ≤ *λ*_*i*_ ≤ *δ*,

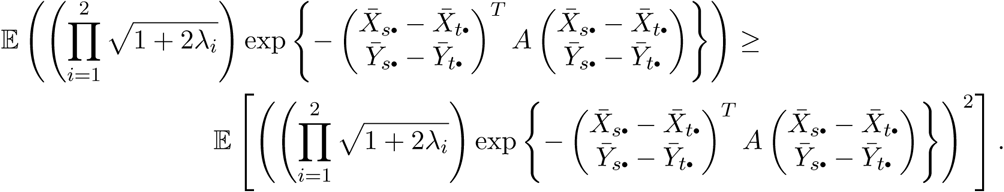

See Appendix Section S1.5 for the proof. In the case where the expectation of an estimator dominates over the variance, reducing the bias tends to be more effective than reducing the variance and keeping a large bias. Using the corrected density estimator, the bias of density term in equation (7) is zero, 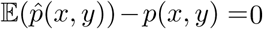. With a zero bias and a small variance, the error in mutual information estimation is also small.

The condition of 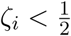 in Theorem 2 and small *λ_i_* in Claim 1 may not hold with a predefined bandwidth. We take a heuristic strategy of shrinking the error covariance by a scalar and using the shrunk covariance to construct the estimator. Using this strategy, the corrected density estimator is no longer asymptotically unbiased. Nevertheless, the corrected estimator with the shrinking covariance strategy still performs better than the baseline that directly uses 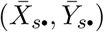 in the KDE (5) and mutual information estimation (6), as we show in Results Section.

### 2.4 Relaxing the assumption of the same error distribution across samples

Biological datasets usually do not have the same distribution of measurement errors across all samples or datasets. Even within datasets, the error distribution varies because of the domain of the measurements or the constraints of the computational methods. For example, gene expression is constrained to be nonnegative. Thus the standard deviation of lowly expressed genes is lower compared to that of highly expressed genes. Therefore, we adapt our corrected estimators for the case where different samples have different measurement errors.

In the semi-discrete case, the error distribution assumption can be relaxed so that each joint category has its unique error distribution. Under the relaxed error distribution assumption, the entry in transition matrix *F*_*ji*_ can be calculated using the error distribution for category *C*_*j*_. The corrected PMF 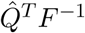 is still an asymptotically unbiased estimator.

In the continuous case using KDE, the error distribution assumption can be relaxed so that each sample has a unique error distribution. Each KDE term 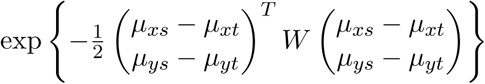 can be corrected using the sum of error covariance in sample *s* and *t*. However, the estimated error covariance may be less accurate because there are only *B* bootstraps to estimate the sample-specific error distribution, compared to estimating the covariance for all samples from *SB* bootstraps.

## 3 Results

### 3.1 Corrected PMF leads to more accurate mutual information estimation in simulated semi-discrete signals

We simulate the semi-discrete signals by the following procedure. Each of *ξ*_*x*_ and *ξ*_*y*_ have 5 categories with real-valued measurements corresponding to [*i* − 1, *i*) for 1≤ *i* ≤ 5. A ground truth PMF is simulated from the 5 × 5 categories with a Dirichlet prior. The true signals within each category follows a uniform distribution in the space of [*i* − 1, *i*) × [*j* − 1, *j*). Each joint category has its unique measurement error distribution, which is a mixture of two Gaussian distributions. Using these probability distributions, we simulate observations using various numbers of samples (100, 500, 1000, 10 000 and 100 000) and various numbers of bootstrap measurements (10, 20, 50 and 100). We compare our corrected estimator with the baseline estimator that treats 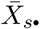 and 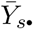 as true signals for estimating PMF and mutual information.

With a fixed bootstrap size and a large sample size, our correction in more accurate in estimating PMF as well as mutual information (Figure 2A, C). Specifically, when the sample size is larger than or equal to 500, the corrected estimator shows its improvement compared to the baseline in both PMF and mutual information estimation. As the sample size grows, the improvement becomes more noticeable. With a small sample size, the sample size is the bottleneck of accurately estimating the PMF, and therefore the corrected estimator does not show an improvement.

**Figure 2:**
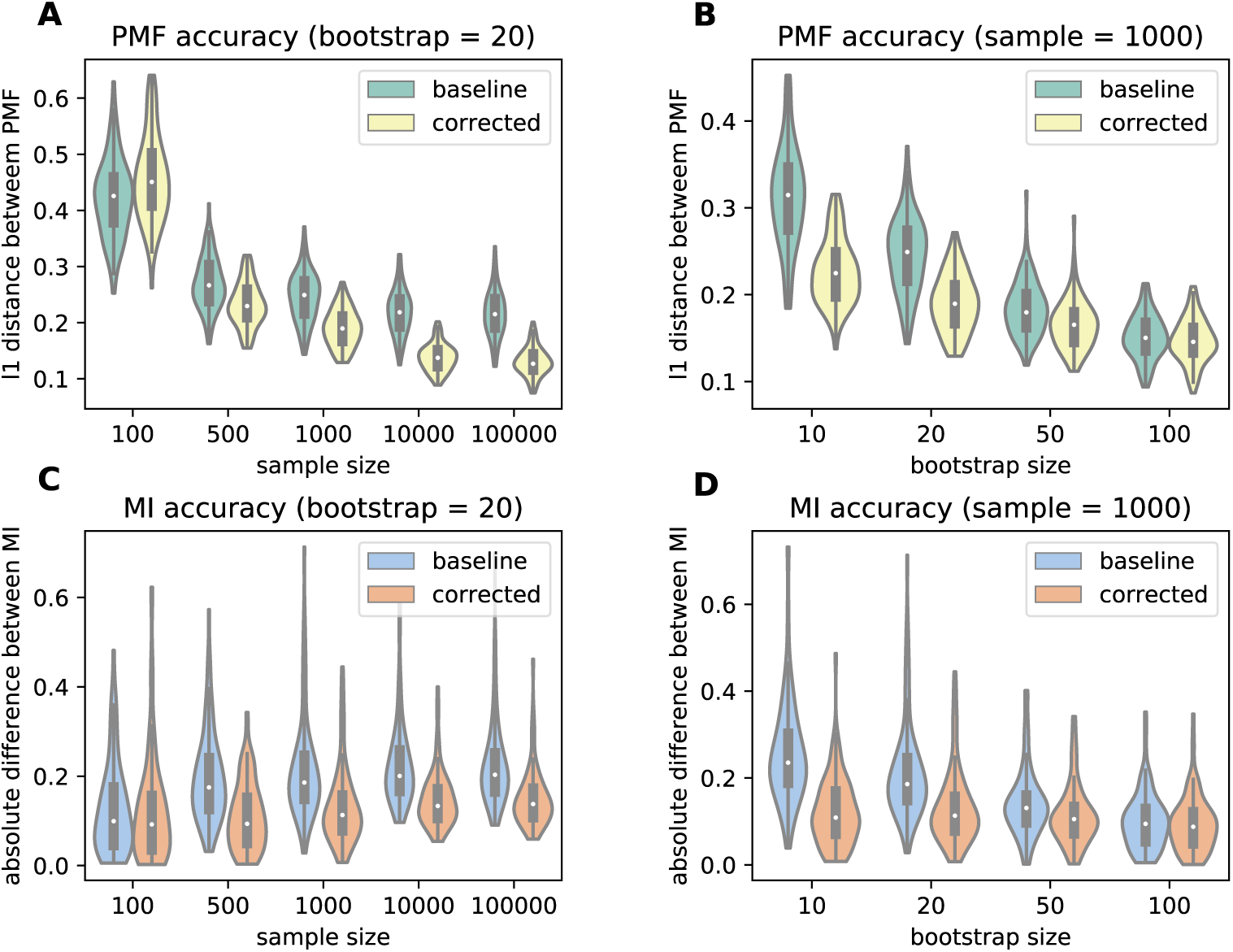
(A–B) *ℓ*_1_ distance between the true PMF and estimated PMF when using baseline (average of bootstraps) estimator and the corrected estimator. (A) The number of bootstraps is fixed to be 20 and the number of samples are indicated by x axis. (B) The number of samples is fixed to be 1000 and the number of bootstraps are indicated by x axis. (C–D) Absolute difference between true mutual information and estimated mutual information when using baseline PMF and corrected PMF. (C) The number of bootstraps is fixed to be 20 and the number of samples is indicated by x axis. (D) The number of samples is fixed to be 1000 and the number of bootstraps is indicated by x axis.

Fixing the bootstrap size, the estimation error for mutual information increases as the number of samples increases for both corrected and baseline estimator (Figure 2C). The baseline PMF estimation converges to a different distribution from 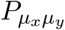 and thus is more biased, which explains its increasing error. The estimation error of the corrected estimator increases more slowly than the baseline. However, it does not converge to the true mutual information either, which is possibly because the error is non-Gaussian and suffers from estimation error with a fixed bootstrap size.

With a fixed sample size, increasing the number of bootstraps reduces the estimation error for both baseline and corrected estimator (Figure 2B, D). This can be explained by that both the baseline and the corrected estimator are asymptotically unbiased as the number of bootstraps goes to infinity. However, before the number of bootstraps is sufficiently large, the corrected estimator achieves a smaller estimation error.

### 3.2 Corrected KDE leads to more accurate mutual information estimation for simulated Gaussian mixtures

The true signal (*µ*_*x*_, *µ*_*y*_) is simulated from a mixture of bi-variated Gaussian distributions. In each of the mixture, the measurement error is a mixture of two Gaussian distributions that have smaller covariances than the Gaussian covariance matrix of (*µ*_*x*_, *µ*_*y*_). We simulate various numbers of mixtures (2, 5, 10 and 20), various numbers of samples (500, 1000, 5000 and 10 000), and various numbers of bootstraps (20, 50 and 100). Since there is no closed-form expression of mutual information for a mixture of Gaussian distributions, we apply the KDE-based mutual information estimation (6) on the simulated true signals (*µ*_*xs*_, *µ*_*ys*_) and use it as the true mutual information. When calculating the corrected estimation in equation (10), we split the samples into clusters by applying *K*-means clustering on the sample-specific error covariance and estimate an error covariance using all samples in each cluster. The number of *K*-means clusters is set to the same as the number of Gaussian mixtures. The error covariance matrix is shrunk so that *ζ*_*i*_ < 0.25 to satisfy the conditions in Theorem 3 and Claim 1. We compare to a baseline estimator that uses the average of bootstraps in the KDE-based mutual information estimator in (5) and (6). The bandwidths are set to be 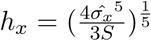 and 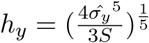 as suggested by Silverman [22] for both estimators.

With a fixed number of bootstraps, we observe the same pattern as in the semi-discrete case: the corrected mutual information estimator achieves smaller biases than the baseline (Figure 3A). When the sample size becomes larger, the improvement of the corrected estimator becomes more apparent because the estimated density converges to its expectation.

**Figure 3:**
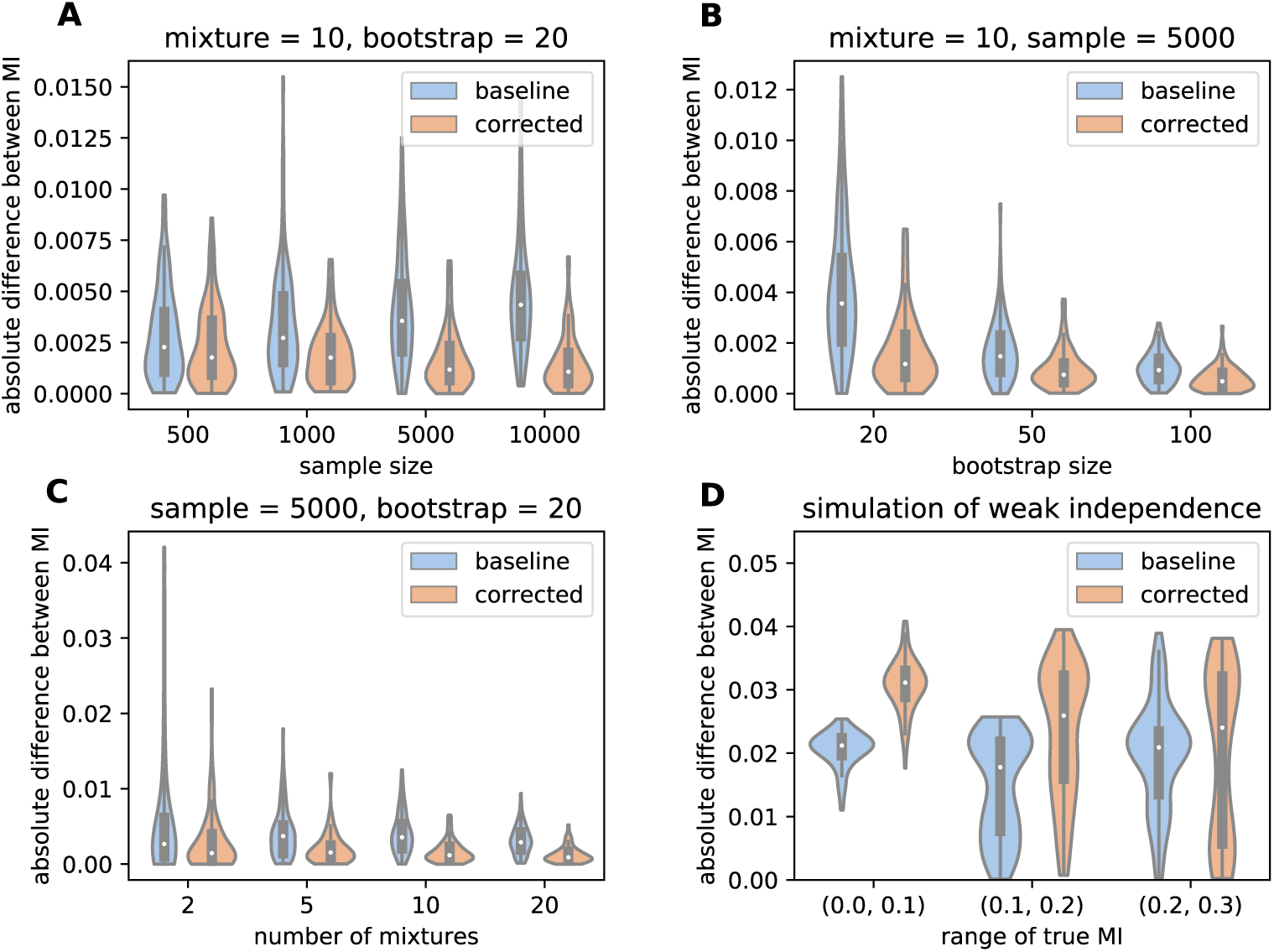
Absolute difference between the true mutual information and estimated mutual information using the baseline estimator and the corrected estimator. (A) The mixture size is 10, the bootstrap size is 20, and the sample size is indicated by x axis. (B) The mixture size is 10, the sample size is 5000, and the bootstrap size is indicated by x axis. (C) The sample size is 5000, bootstrap size is 20, and the mixture size is indicated by x axis. (D) Simulation of weakly dependent random variables. The true signals follows a bi-variate Gaussian distribution. The sample size is 5000 and the bootstrap size is 20. X axis is the true mutual information.

With a fixed number of samples, increasing the number of bootstraps reduces the mutual information estimation error for both estimators (Figure 3B). The improvement of the corrected estimator is visible even with 100 bootstraps, especially in terms of the smaller variance of estimator error.

We observe that the correction is effective for various numbers of mixtures (Figure 3C), but the effectiveness is more significant under a larger number of mixtures. Using Mann-Whitney one-sided U test to compare the accuracy between the baseline and corrected mutual information, the p-value is 0.002 for two mixtures, 1.41×10^−7^ for five mixtures, and 4.46^−16^ for ten mixtures under 5000 samples and 20 bootstraps. Nevertheless, a wider range of mixture sizes needs to be tested in order to study the effect of the number of mixtures on mutual information estimation in more detail.

The corrected estimator is able to reveal strong dependencies when they are shadowed by the measurement error. However, when the dependence is weak or even the two signals are independent, the correction may lift the estimated mutual information and falsely show a small dependence (Figure 3D). Therefore, the corrected mutual information estimator should be only applied when the dependence is large after correction.

### 3.3 Genes with largest mutual information with known cancer genes are slightly changed by corrected estimator

We applied the corrected mutual information estimator on 1168 breast cancer samples from TCGA to investigate what genes have high mutual information with known cancer genes. TCGA RNA-seq samples are quantified by Salmon [17] with 100 bootstraps for evaluating the measurement error. With 10 arbitrarily chosen cancer genes from COSMIC [30] (cancer.sanger.ac.uk), we estimate the mutual information using both the original KDE-based estimator (uncorrected) with the default single-point Salmon quantification and our corrected mutual information.

In general, the uncorrected and corrected estimator agree with each other. For each of the selected cancer gene, we compute the corrected and uncorrected mutual information between this gene and all other genes. The Spearman correlation between uncorrected and corrected mutual information is greater than 0.98. However, a detailed analysis of the top genes with high mutual information reveals the difference between the two estimators.

Among the 500 genes with highest uncorrected mutual information values for each selected cancer gene, the corrected estimations between the selected genes and those 500 genes vary as much as 10%–40% from the uncorrected ones (Figure 4A, the range of the largest y axis values across selected genes shown on x axis). The sets of genes with the highest estimated mutual information between the two estimators are different when fixing the set size (Figure 4B). Therefore, when the specific values of mutual information are used or when the subset of genes with the highest mutual information is of interest, the measurement error has an effect on the result and the corrected mutual information estimator should be considered.

**Figure 4:**
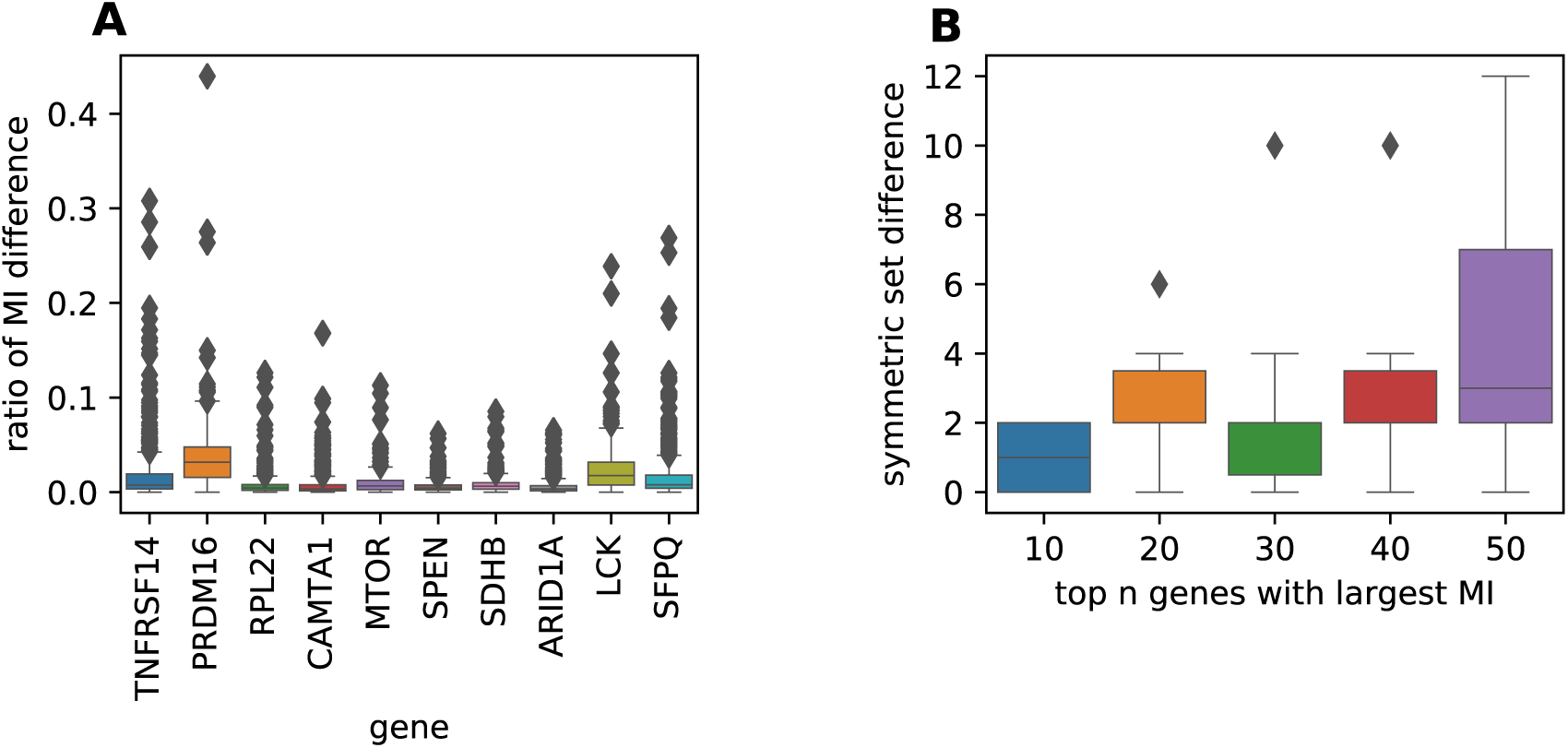
(A) The ratio of change from the uncorrected mutual information estimate to the corrected estimate for the top 500 genes with the largest uncorrected estimates. X axis is the 10 known cancer genes and the box is plotted for the 500 genes with the largest uncorrected mutual information. (B) The symmetric difference between the set of top several genes with largest uncorrected mutual information and the set of top genes with the largest corrected mutual information. X axis is the size of each set and the box is plotted for 10 known cancer genes.

We show two example genes where the corrected estimator decreases its ranking of mutual information. The rank of estimated mutual information between gene *CASP9* and known cancer gene *SDHB* decreases from 91 to 67 (the estimate changes from 0.198 to 0.211). *SDHB* affects mitochondrial function and *CASP9* helps cell apoptosis [31]. Mitochondria plays a role in cell apoptosis [32], providing support that the expressions of the two genes possibly are dependent. The rank of the estimate between genes *GRAP2* and *LCK* decreases from 86 to 71 (the estimate changed from 0.483 to 0.509). Both *GRAP2* and *LCK* are involved in T-cell-receptor signaling pathway [33].

In this analysis, there is no ground truth about the true mutual information or the ranking of the pairwise mutual information. In addition, the number of biological samples may not be large enough to reveal the true dependence between genes. A larger number of samples and a better validation are needed for a more comprehensive evaluation of the performances of the corrected and uncorrected estimator.

## 4 Discussion

We derive a corrected mutual information estimator to account for the measurement error for both semi-discrete and continuous measurements. Our corrected estimator is based on estimating the probability mass function (PMF) or probability density function (PDF) in an asymptotically unbiased way. We prove that in the semi-discrete case the corrected mutual information estimator derived from the unbiased PMF estimation is asymptotically unbiased. We give conditions under which our corrected estimator in the continuous case reduces the bias of mutual information estimation. On simulated data, the corrected estimators for both semi-discrete and continuous cases are more accurate compared to the baseline of using the bootstrap average in mutual information estimation.

We compare our corrected estimator to the uncorrected estimator on detecting the genes with high dependence with known cancer-related genes using TCGA breast cancer gene expression data. The estimated mutual information is generally consistent between the corrected and uncorrected estimator. However, the values of the estimated mutual information may change up to 40% for the top 500 dependent genes. The sets of a fixed number of top dependent genes differ by a few number of genes between the two estimators. The observation suggests that the measurement error has an effect on the mutual information estimation and should be accounted for when carrying out analyses based on the value or ranking of estimated mutual information.

Our corrected estimator for continuous random variables reduces the bias of mutual information estimation only under certain assumptions. Whether there is an unbiased estimator and the form of the estimator remains to be determined. In addition, a correction for the KNN-based mutual information estimator to correct for measurement error is also a useful future direction.

Our corrected estimators assume that the measurement error has zero mean and does not have systematic biases. However, this assumption is unrealistic for certain data types. Modeling and correcting the systematic biases is also critical for accurately estimating the mutual information, and this is an important direction for future work.

We focus on reducing the bias of mutual information estimation. However, low variance or other properties may also be desired. Designing new corrections with the consideration of measurement error and studying their statistical properties will be interesting theoretical future directions.

## Acknowledgements

This research is funded in part by the Gordon and Betty Moore Foundation’s Data-Driven Discovery Initiative through Grant GBMF4554 to C.K., by the US National Science Foundation (CCF-1256087, CCF1319998) and by the US National Institutes of Health (R01GM122935). This work was partially funded by The Shurl and Kay Curci Foundation. This project is funded, in part, under a grant (#4100070287) with the Pennsylvania Department of Health. The Department specifically disclaims responsibility for any analyses, interpretations or conclusions. The results shown here are in part based upon data generated by the TCGA Research Network: https://www.cancer.gov/tcga. This work used the Extreme Science and Engineering Discovery Environment (XSEDE), which is supported by National Science Foundation grant number ACI1548562. Specifically, it used the Bridges system, which is supported by NSF award number ACI-1445606, at the Pittsburgh Supercomputing Center (PSC) [34]. The results shown here are in whole or part based upon data generated by the TCGA Research Network: https://www.cancer.gov/tcga.

We thank Guillaume Marçais, Hongyu Zheng, and Amir Alavi for useful suggestions.

## S1 Proofs

### S1.1 The corrected estimator in semi-discrete case is unbiased in estimating entropy

Let 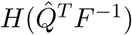 be the entropy calculated using PMF 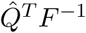 and similarly *H*(*Q*^*T*^ *F*^−1^) be the entropy calculated using *Q*^*T*^ *F*^−1^. Let *G* = *F* ^−1^ and *g*_*ij*_ be the entry of *G* on the *i*^*th*^ row *j*^*th*^ column.

#### Lemma S2.

*Let Q be the underlying PMF of g*(*X*_*s*1_, *Y*_*s*1_). *Given S samples of* (*X*_*s*1_, *Y*_*s*1_). *Let* 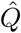 *be the estimator of Q given by*

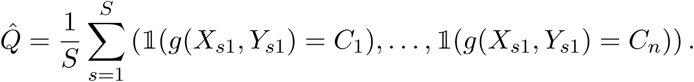

*Given matrix G* ∈ ℝ^*n*×*n*^, *the entropy calculated using* 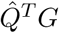 *is an unbiased estimator of the entropy calculated using Q*^*T*^*G as S* → ∞

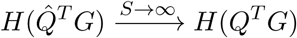

*Proof*. We use the proof from Basharin [20] as a template. Basharin [20] derived the Taylor expansion of the entropy:

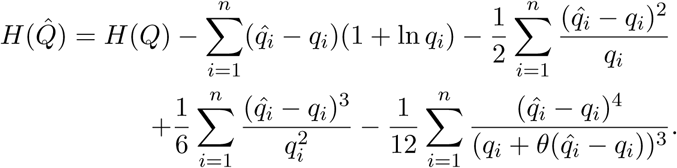

They derive the following asymptotic values of the expectation of each term in the above equation:

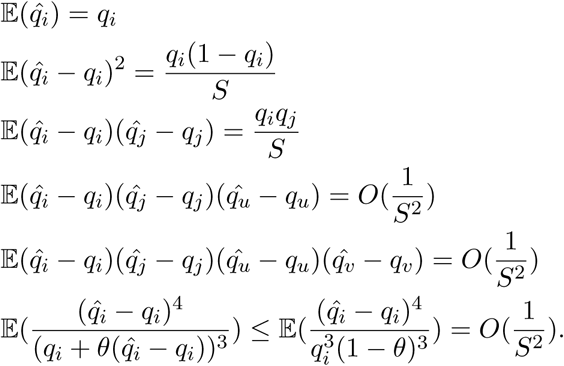

Using their derivation, the Taylor expansion of 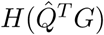 is

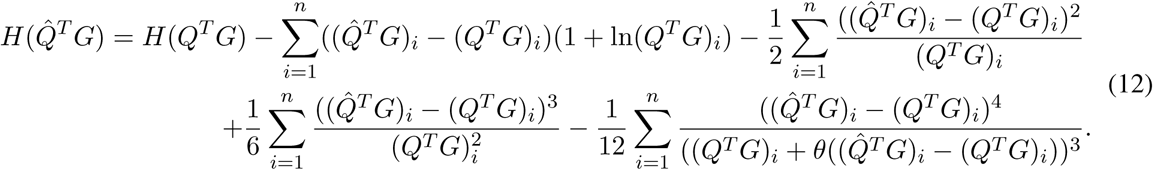

The term 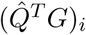 and (*Q*^*T*^*G*)_*i*_ can be explicitly expressed as 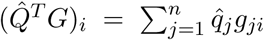 and 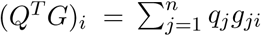. Using the explicit expression, the expectation of the terms in the Taylor expression is

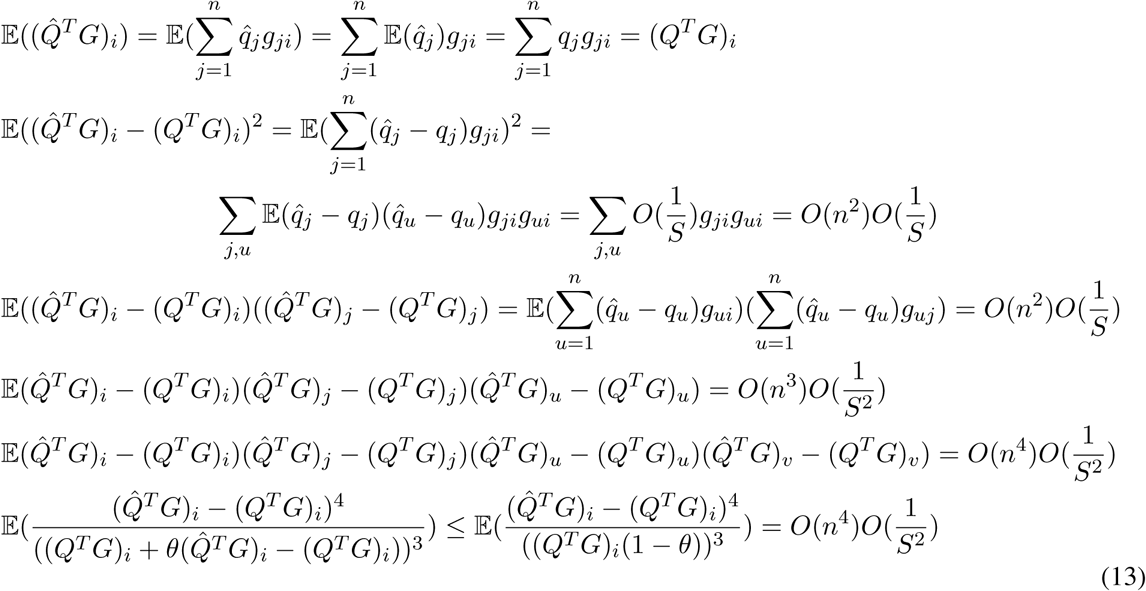

Plugging the equations and inequalities in (13) in the expectation of equation (12), the expectation of the entropy estimator is

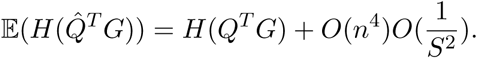

Since the number of categories *n* is fixed and the number of samples *S* goes to infinity,

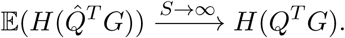

Hence, we proved that the entropy 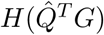 is an unbiased estimator for entropy *H*(*Q*^*T*^*G*) as the number of samples *S* goes to infinity.

### S1.2 Estimation bias of log term using estimated density 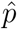

#### Lemma 1.

*Given the true probability density p and fixed point* (*x, y*), *for any estimator of the density at the point* 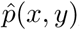, *if the true and estimated density are lower bounded by δ* > 0, *that is*, *p*(*x, y*) ≥ *δ and* 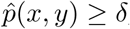, *then the bias of* 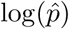 *at the point is upper bounded by*:

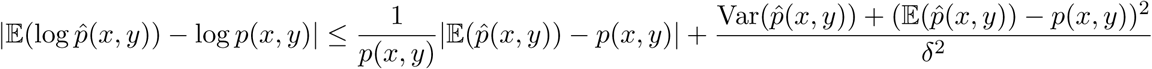

*Proof*. The Taylor expansion on the log function with the mean value form states that there exists *θ* ∈ (0, 1) such that

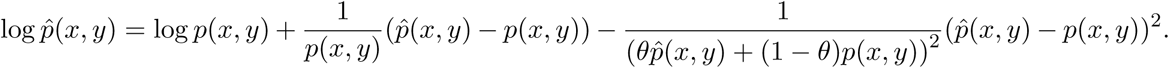

Using the lower bound *δ* of *p*(*x, y*) and 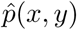, the second-order term can be bounded by

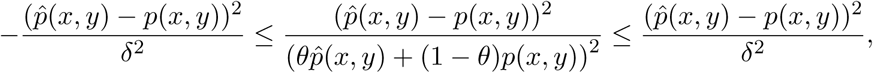

and thus the following inequalities hold:

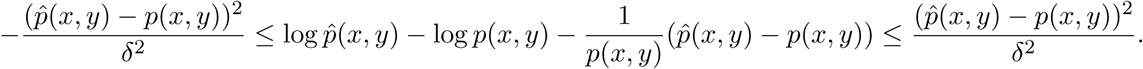

Taking the expectation on both sides, the equation can be arranged in the following way:

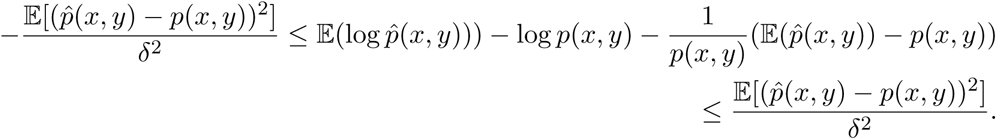

The two inequalities can be combined using the absolute value form:

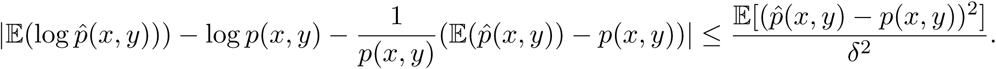

Applying the triangle inequality of the absolute value on the left hand side, we have:

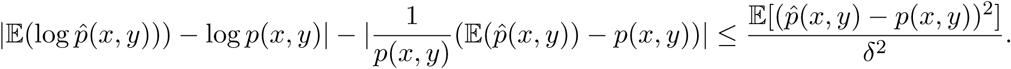

Replacing the term 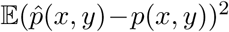 with the bias-variance decomposition formula, therefore, we derive the upper bound of the estimation bias of the log term:

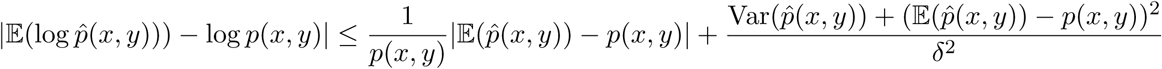

### S1.3 The corrected density estimator in the continuous case is unbiased in estimating the exponential term in KDE

We use the following lemma to prove the unbiasedness of density estimator.

#### Lemma S3.

*Given n-dimensional Gaussian random variable X* ~ *N* (*µ*, Σ) *and matrix A* ∈ ℝ^*n*×*n*^, *the moment generating function for Q* = *X*^*T*^*AX* is

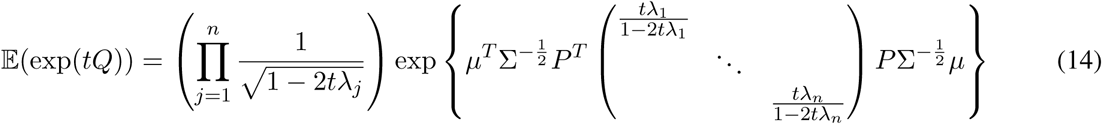

*where P and λ*_*i*_ *are the eigenvectors and eigenvalues of* 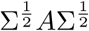, that is, 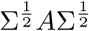, *and t is constrained by* 1 − 2*tλ*_*i*_ > 0 *for all i*.

*Proof*. The proof was originally given by user “kjetil b halvorsen” on StackExchange (https://stats.stackexchange.com/questions/262604/what-is-the-moment-generating-function-of-the-generalizedmultivariate-chi-squ/318908#318908). We reproduce it here for completeness to make our argument self contained.

*P* and *λ*_*i*_ are the eigenvectors and eigenvalues of 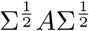, they can be equivalently expressed by the equation of 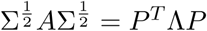, where *P* is orthonormal and Λ = diag(*λ*_1_, *λ*_2_, …, *λ*_*n*_).

Let 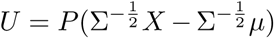 and 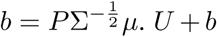. *U* + *b* is an affine function of Gaussian random variable *X*, thus the distribution of *U* + *b* is a also Gaussian distribution. The expectation is

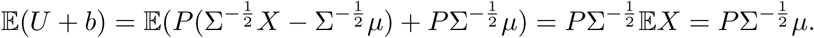

The covariance matrix is

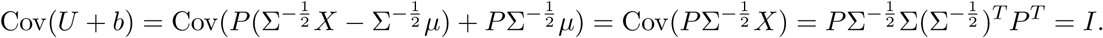

Therefore, *U* + *b* follows a Gaussian distribution where each entry is independent of the others:

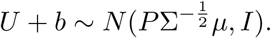

Connecting *U* + *b* back to *Q*, one can verify that 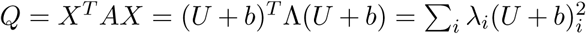, which is the weighted sum of *n* independent non-central chi-squared random variables. Therefore,

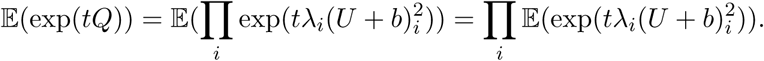

The product can be moved out of the expectation due to the independence between the chi-squared random variables (*U* + *b*)_*i*_. Each of the summation term is the moment generating function of the non-central chi-squared random variable evaluated at point *tλ*_*i*_. Let 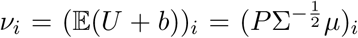. When 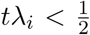, the moment generating function evaluated at *tλ*_*i*_ is

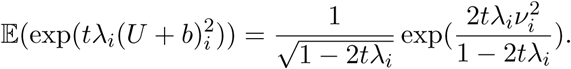

When the inequality 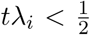 holds for all *i*, all the moment generating functions is finite at the corresponding point, and all terms in the production is well-defined. Plugging the expression of each term in the production,

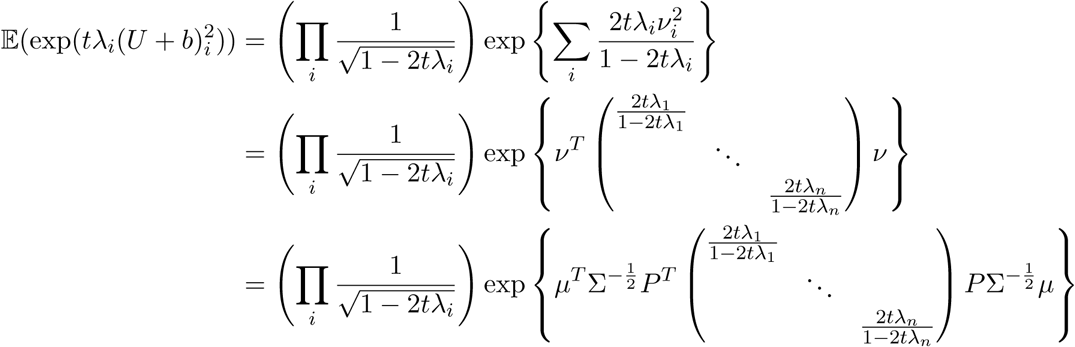

#### Theorem 2.

*Let* 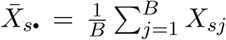 *and* 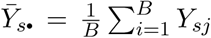. *Let W be an diagonal bandwidth matrix used in KDE. Assuming the measurement error of* 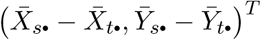 *follows a Gaussian distribution* 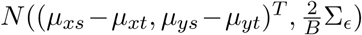, *let P and* {*ζ*_1_, *ζ*_2_} *be the eigenvectors and eigenvalues of* 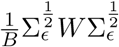. *Let* 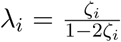, 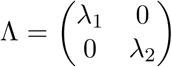, *and* 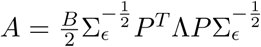. *When ζ*_*i*_ *satisfies* 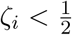 *for all i, the following estimator*,

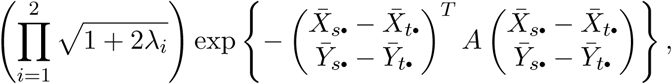

*is an unbiased estimator for the KDE term in* *equation* (5)

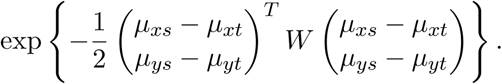

*Proof*. Plugging in *t* = −1 and 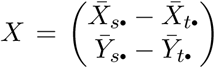 in Lemma S3. After calculating the eigenvalues and eigenvectors of 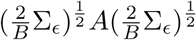, we have the eigenvalues are {*λ*_1_, *λ*_2_} and the eigenvectors are *P*. When expressing *ζ*_*i*_ using *λ*_*i*_, the constraint 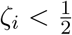 under *t* = −1 indicates that 1 − 2*tλ*_*i*_ > 0, that is, the condition of the above lemma holds. Thus, the expectation of the estimator is:

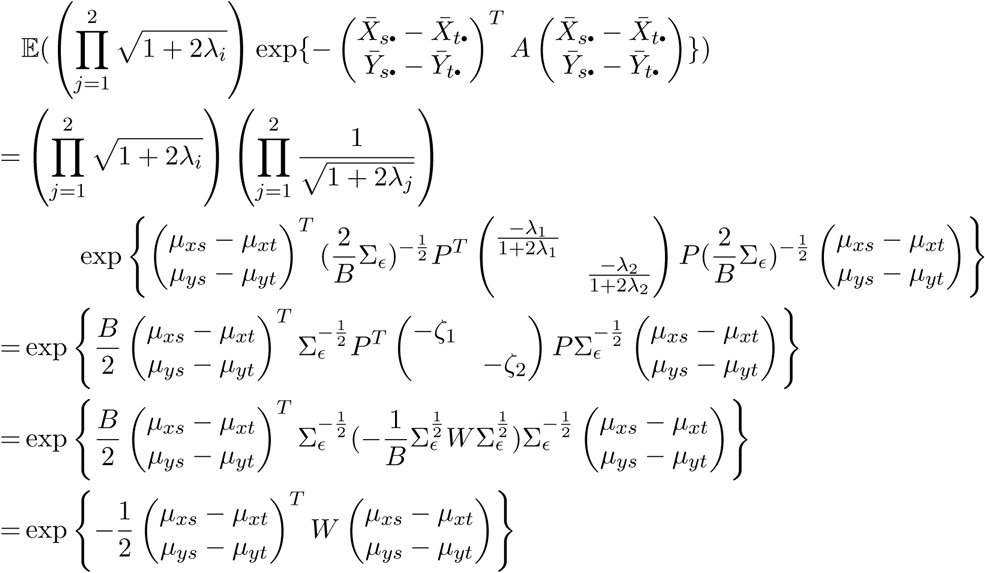

### S1.4 Corrected density estimator is asymptotically unbiased when using estimated error covariance

#### Lemma S4.

*Given a series of n-dimensional Gaussian random variables* 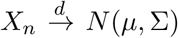 *where each of X*_*n*_ *has a finite moment generating function for t* ∈ (*a, b*). *Given matrix A* ∈ ℝ^*n*×*n*^, *the moment generating function for* 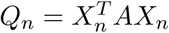 is

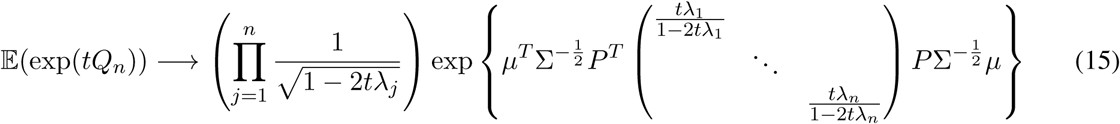

*pointwise for each t values within the region* 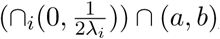, *where P and λ*_*i*_ *are the eigenvectors and eigenvalues of* 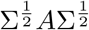.

*Proof*. By the same construction of 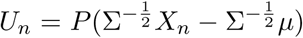 and 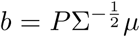, the limiting distribution is also a Gaussian distribution by an affine transformation of the random variable:

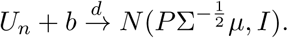

The limiting distribution for 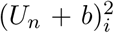 is a chi-squared distribution because of the following. Let 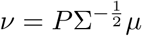. The cumulative distribution function of 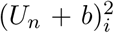 is:

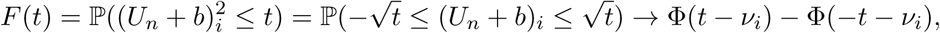

where the right hand side is the same as the cumulative distribution function of a non-central chi-squared distribution with degrees of freedom equal to 1.

Chareka [35] has proved that when the moment generating function of each (*U*_*n*_ + *b*)_*i*_ is finite at *t* ∈ (*a, b*), the convergence of distribution implies the convergence of moment generating function. Using the moment generating function calculated in Lemma S3, 𝔼(exp(*tQ*_*n*_)) converges to the moment generating function of the summation of chi-squared random variables in the limiting distribution.

#### Theorem 3.

*Let P and* {*ζ*_1_, *ζ*_2_} *be the eigenvectors and eigenvalues of* 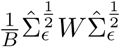. *Let* 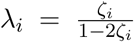, 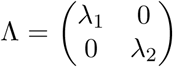, *and* 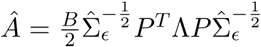. *When ζ*_*i*_ satisfies 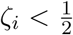 *for all i, the following estimator is an asymptotically unbiased estimator for the KDE term in* *equation* (5):

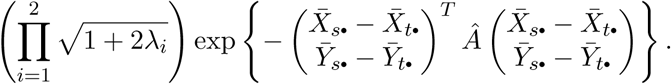

*Proof*. The proof is almost the same as the proof for Theorem 2 except that Lemma S4 is used for calculating the moment generating function of the limiting distribution. Set *t* = −1 and 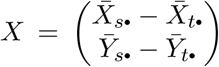. The expectation of the estimator approaches to

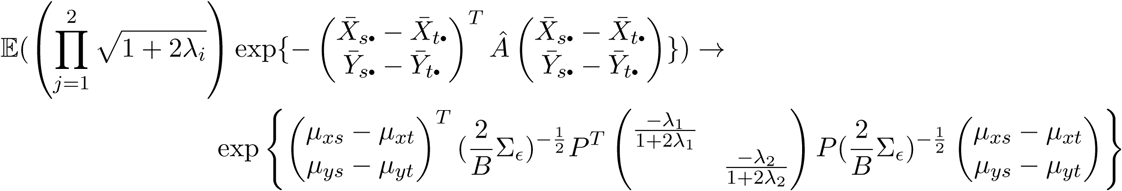

Substituting the value of *λ*_*i*_ and the equality relationship between *P*, Σ_*ϵ*_, and *ζ*_*i*_, the above expectation converges to the KDE term of the true signal values:

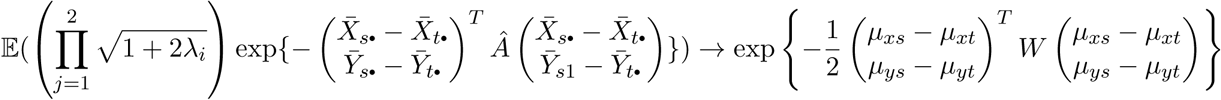

### S1.5 The expectation of corrected density estimator is larger than its second order moment when *λ*_*i*_ is small

**Claim 1.** *There exists a small positive number δ such that when* 0 ≤ *λ*_*i*_ ≤ *δ*,

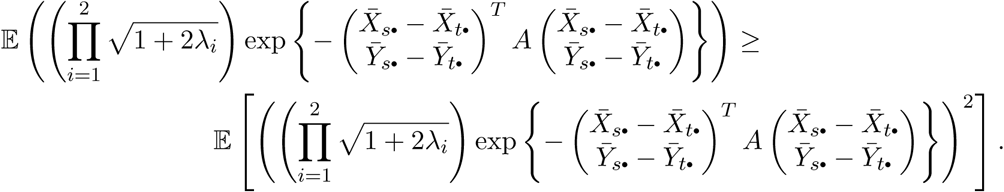

*Proof*. The expectation, as the first step of proof of Theorem 2, is

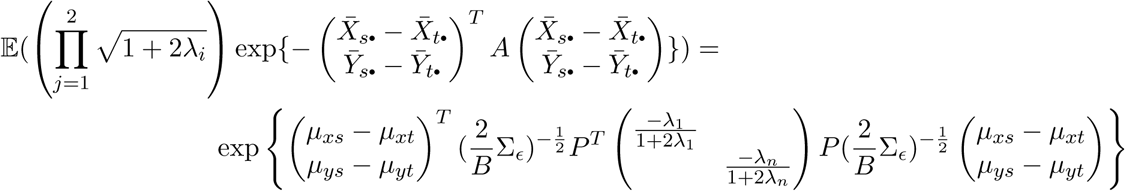

The second order moment around 0 has the expression of

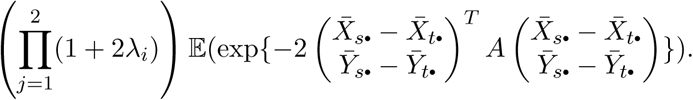

This can be calculated by the same steps as in the proof of Theorem 2 except setting *t* = −2:

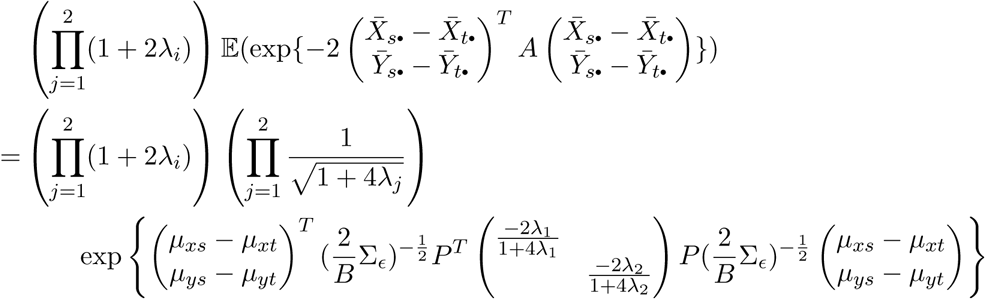

Let 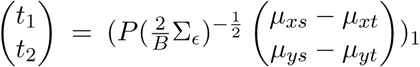. The ratio between the expectation and the second order moment is

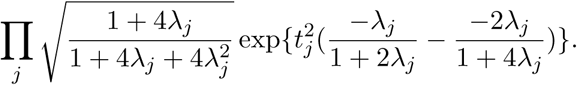

We need to prove that within a small positive region (0, *δ*], each product term in the above equation is greater than 1. Or equivalently, we need to prove the rearranged form:

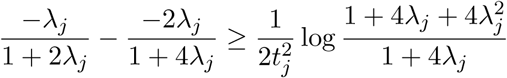

Let function 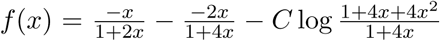, where 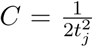. The function is 0 when *x* takes value of 0, that is, *f* (0) = 0. When *x* > 0, the derivative is

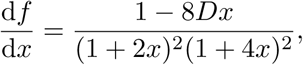

where *D* = 1 + *C*(1 + 4*x*)(1 + 2*x*). Thus, there exists a positive region (0, *δ*] such that when *x* ∈ (0, *δ*], 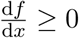. And in this region,

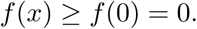

This is equivalent to:

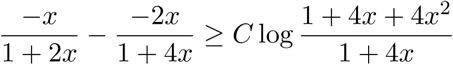

Therefore, when *λ*_1_, *λ*_2_ ∈ (0, *δ*], the ratio between the expectation and the second order moment is larger than 1, and the second order moment is upper bounded by the expectation.

